# Hyperthermia reduces antimicrobial tolerance of Mycobacterium marinum in biofilms and ex vivo zebrafish granulomas

**DOI:** 10.1101/2025.11.28.691154

**Authors:** Alina Jokinen, Milka Hammarén, Rosa Korhonen, Mataleena Parikka

**Affiliations:** Faculty of Medicine and Health Technology, Tampere University, Tampere, Finland; European Molecular Biology Laboratory, Heidelberg, Germany

**Author notes:** Equal contribution (Alina Jokinen, Milka Hammarén).

**Keywords:** Mycobacterial biofilms, antimicrobial tolerance, hyperthermia, tuberculosis, heat therapy, antimicrobial persistence, Mycobacterium marinum

## Abstract

The treatment of tuberculosis (TB) is slow and inefficient. One suggested reason for the treatment difficulties are drug-tolerant biofilms that have been shown to develop in TB patients and animal models. In this pioneering pilot study, we explored the potential of hyperthermia in sensitizing biofilm-associated tolerant mycobacteria to antimicrobial agents. As the starting point, we used our newly-developed in vitro biofilm minimum duration for killing (MDK) tolerance assay for screening hyperthermal treatments of bioluminescent mycobacteria. Using standard plating methods, we verify that the combination of hyperthermia at 47°C for 30 minutes and an antimicrobial drug rifampicin at 400 μg/ml has positive effects against cultured *Mycobacterium marinum* biofilms as well as *M. marinum* from granulomas isolated from adult zebrafish with chronic-stage infection. These intriguing findings suggest that hyperthermia could be used to enhance conventional TB treatments.

**Author Summary:** Tuberculosis (TB) and similar infections are hard to cure because the bacteria can “hide” from antibiotics. They do this by entering a state where they tolerate drugs without becoming resistant, often forming biofilms—sticky clusters that protect them from treatment. This makes therapy long and relapse common. We asked whether heat could help break this tolerance. Using a new lab test, we combined heat at different durations and temperatures with antibiotics against *Mycobacterium marinum*, a close relative of the TB bacterium. Heating made the antibiotic rifampicin much more effective, both in biofilms and in infected tissue from zebrafish, which mimic TB-like lesions. We saw similar benefits in TB bacteria when heat was paired with streptomycin. These results suggest that controlled heating could work alongside standard drugs to kill stubborn bacteria faster. Because heat damages cells in a general way, not by targeting one molecule, it might also reduce the risk of antibiotic resistance

## Introduction

Tuberculosis (TB) and other mycobacterial infections remain notoriously hard to treat, largely because of antimicrobial tolerance and drug resistance. Resistance allows bacteria to grow despite antimicrobials, whereas tolerance simply lets them endure prolonged exposure without acquiring genetic resistance (1). This silent tolerance matters: tolerant populations often create the conditions for resistance to emerge, making tolerance an important target in its own right (2). For a variety of bacterial species it has been shown that antimicrobial tolerance is increased in biofilms; structured communities embedded in an extracellular matrix that shield bacteria from immune attack and contribute to chronic infection as well as drug tolerance (3). For years, mycobacterial biofilms have been known to exhibit strong tolerance in vitro (4–6). Within the biofilm microenvironment, bacteria are protected by an extracellular polymeric substance (EPS) matrix, reduced metabolic activity, and altered microenvironments, all of which collectively impair antibiotic penetration and efficacy (7,8). Only recently, however, have similar biofilm structures been observed in vivo, underscoring their clinical relevance (9). This discovery may help explain why current treatments are so lengthy, complex, and prone to relapse (3). Novel therapeutic strategies are urgently needed to enhance bacterial clearance and prevent recurrence.Tuberculosis (TB) and other mycobacterial infections remain notoriously hard to treat, largely because of antimicrobial tolerance and drug resistance. Resistance allows bacteria to grow despite antimicrobials, whereas tolerance simply allows them to endure prolonged exposure without acquiring genetic resistance (1). This silent tolerance matters: tolerant populations often create the conditions for resistance to emerge, making tolerance an important target in its own right (2). For a variety of bacterial species it has been shown that antimicrobial tolerance is increased in biofilms; structured communities embedded in an extracellular matrix that shield bacteria from immune attack and contribute to chronic infection as well as drug tolerance (Ciofu et al., 2022). For years, mycobacterial biofilms have been known to exhibit strong tolerance in vitro (3–5). Within the biofilm microenvironment, bacteria are protected by an extracellular polymeric substance (EPS) matrix, reduced metabolic activity, and altered microenvironments, all of which collectively impair antibiotic penetration and efficacy (6,7). Only recently, however, have similar biofilm structures been observed in vivo, underscoring their clinical relevance (8). This discovery may help explain why current treatments are so lengthy, complex, and prone to relapse (9). Novel therapeutic strategies are urgently needed to enhance bacterial clearance and prevent recurrence.

Heat-based therapies have recently gained attention as an adjunct to conventional antimicrobials for biofilm eradication, especially in the context of biofilm infections of implants (10). Several studies have demonstrated that mild or transient heating can disrupt biofilm integrity, increase bacterial metabolic activity, and sensitize cells to antimicrobial drugs. For instance, thermal mitigation of *Pseudomonas aeruginosa* biofilms showed that elevated temperature alone can impair biofilm stability (11), while combining heat with antimicrobial agents resulted in marked synergistic effects (12,13). Similar findings have been observed with *Staphylococcus epidermidis* and *Staphylococcus aureus*, where hyperthermia or thermal shock potentiated the bactericidal activity of various antimicrobials (14,15). Importantly, the combination of heat and antimicrobials appears robust across bacterial species, suggesting that thermal stress can overcome multiple tolerance mechanisms. In addition, in the cancer therapy field, various hyperthermic treatments have been successfully used in patients (16–18) suggesting the clinical applicability and feasibility of therapeutic hyperthermia in humans.

The mechanisms underlying these synergistic effects are multifactorial. Heat can increase membrane permeability, enhance diffusion of antibiotics through the biofilm matrix, and transiently stimulate metabolic activity, thereby improving the efficacy of drugs that target active cellular processes (15,19,20). Moreover, approaches such as magnetic nanoparticle-induced hyperthermia (19–21), photothermal nanomaterials (22–25), and alternating magnetic fields (21,26,27) have expanded the potential for localized, non-invasive heating. These technologies not only enable spatial and temporal control over biofilm disruption but also open possibilities for combining physical and pharmacological modalities against multidrug-resistant pathogens (22,28,29). Collectively, these studies highlight that mild heat treatment can act as a powerful antimicrobial adjuvant by shifting the physiological state of biofilm-embedded bacteria.

Despite these advances, most investigations into thermal–antimicrobial combinations have focused on *Staphylococcus* or *Pseudomonas* species, as well as foodborne pathogens such as *Klebsiella pneumoniae* (30). To our knowledge, there are currently no reports on how biofilms formed by slow-growing mycobacteria, including *Mycobacterium tuberculosis* and non-tuberculous mycobacteria (NTM), respond to combined heat and antimicrobial stress. This represents a significant gap, as mycobacterial biofilms are strongly implicated in persistence, antimicrobial tolerance, and chronic infection (6,9,31,32). Understanding whether hyperthermia can sensitize mycobacterial biofilms to antimicrobial agents could therefore provide novel therapeutic opportunities.

In this study, we investigate the effects of heat treatment in combination with antimicrobial drugs on biofilms of *M. marinum* and *M. tuberculosis* H37Ra, surrogate models for biofilms formed by pathogenic mycobacteria. In addition, we test the effect of the combination treatment on bacteria extracted from granulomas developed in *M. marinum* –infected adult zebrafish. The adult zebrafish model has been used in our laboratory since 2010, is well-established in TB research and recapitulates important aspects of the human disease including caseonecrotic granulomas and dormant bacteria (Parikka et al. 2012, Hammarén et al. 2014). We have also recently shown that the mycobacteria in adult zebrafish also show tolerance to treatment (Lehmusvaara, Sillanpää et al. 2025). By systematically examining different temperature–time regimens and antimicrobial drug combinations, we saw that for *M. marinum* biofilms, a 30-minute thermal treatment at 47°C in combination with rifampicin was clearly beneficial *in vitro* and *ex vivo*. In cultured *M. tuberculosis* biofilms, the effect was present with streptomycin, but was less pronounced. Our findings lay the groundwork for new alternative, physical strategies to combat TB and other recalcitrant mycobacterial infections.

## Results

### Hyperthermia potentiates the antimicrobial activity of rifampicin against M. marinum

To evaluate the heat sensitivity of *M. marinum*, the biofilm minimum duration for killing (MDK) assay was conducted using 15-minute hyperthermia across a temperature gradient ranging from physiological to lethal levels. Incubation at 32 °C was used as the no-heating control. Treatments were performed both in the absence and presence of rifampicin (400 μg/ml). In the absence of the antimicrobial agent, a marked reduction in bacterial viability was observed beginning at 51.8 °C, with complete killing achieved at 60 °C. This defined the upper thermal tolerance limit of *M. marinum* under the tested conditions (Fig. 1a&c). When rifampicin was included after heat treatment, a pronounced enhancement in bacterial killing was observed at sub-lethal temperatures (Fig. 1b&d). Specifically, between 47 °C and 51.8 °C, the combination of heat and rifampicin resulted in approximately two-fold reduction in viability compared to rifampicin alone (Fig. 1d). These findings suggest that mild hyperthermia can potentiate the antimicrobial activity of rifampicin.

**Figure 1.**
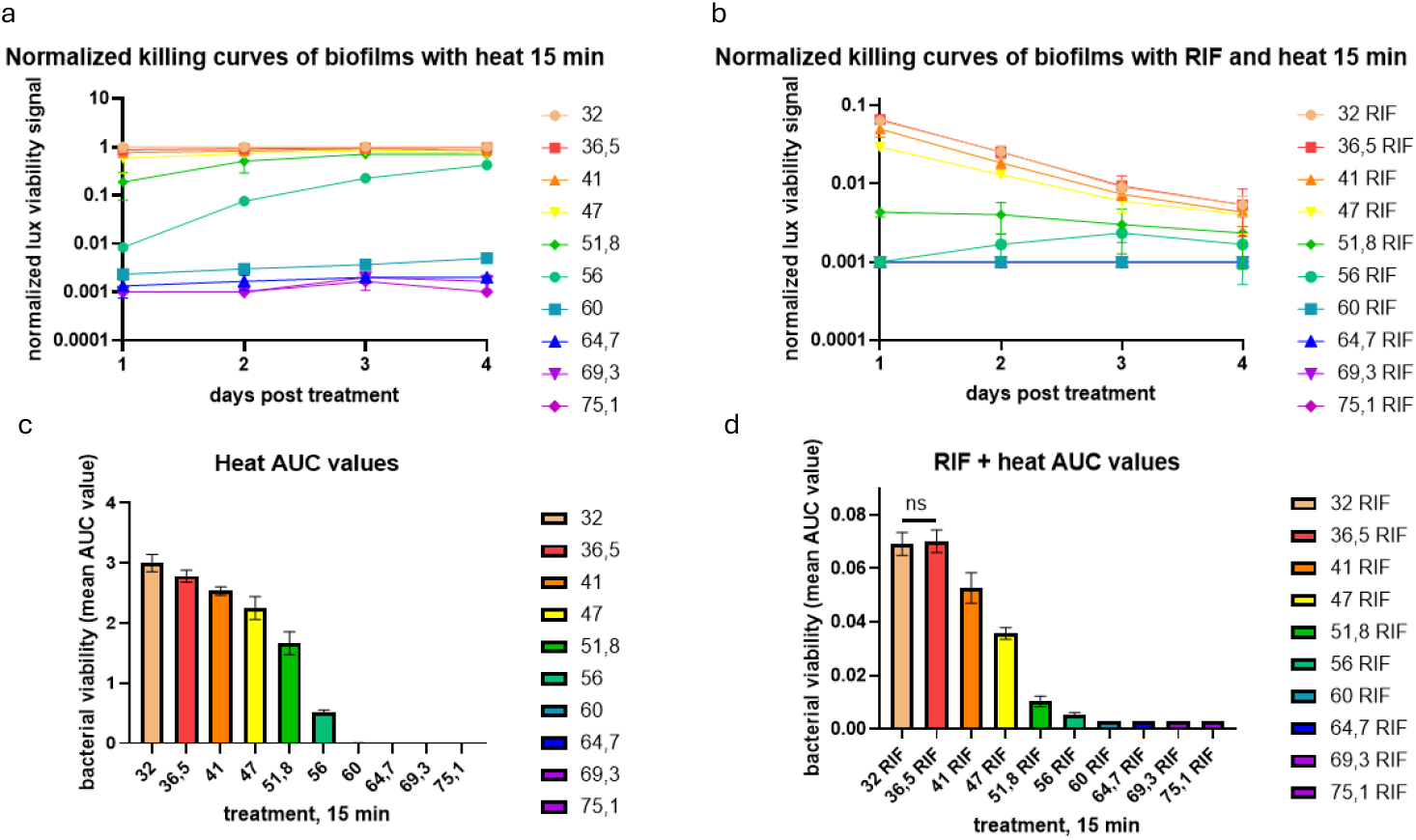
Short hyperthermia potentiates the efficacy of rifampicin against M. marinum biofilms. M. marinum biofilms were exposed to a 15 min heat treatment at increasing temperatures with or without rifampicin. (a) Heat alone reduced viability only at ≥51.8 °C. (b) When combined with rifampicin, sublethal temperatures (47–51.8 °C) markedly enhanced bacterial killing.(c) In the heat-only experiment, the AUC values describing the total effect between 1-4 days after heat treatment were statistically significant compared to 32°C. (d) The AUC values describing the total effect between 1-4 days after heating followed by rifampicin were statistically significant compared to 32°C starting from 41°C (one way ANOVA + Dunnet’s test, n=3).

### Prolonged hyperthermia enhances rifampicin-mediated killing of *M. marinum*

To further investigate the potentiating effect of hyperthermia on antimicrobial tolerance, *M. marinum* samples were exposed to a sub-lethal temperature of 46.8 °C for varying durations (30, 60, 120, and 180 minutes) prior to rifampicin (400 μg/ml) treatment. The bacterial survival was assessed using the biofilm MDK tolerance assay. With increasing duration of treatment, the remaining tolerant population was smaller (Fig. 2a). Colony-forming unit (CFU) analysis performed four days post-treatment showed a marked reduction in bacterial viability across all heating durations when combined with rifampicin, compared to unheated controls (Fig. 2b). Combining a 90-minute heating at 46.8 °C with rifampicin led to full sterilization of the *M. marinum* biofilm culture with no culturable persisters (Fig. 2b) These results demonstrate that a hyperthermal treatment at 46.8°C, when applied for 30-180 minutes, can be beneficial in speeding up killing of tolerant mycobacteria in biofilms.

**Figure 2.**
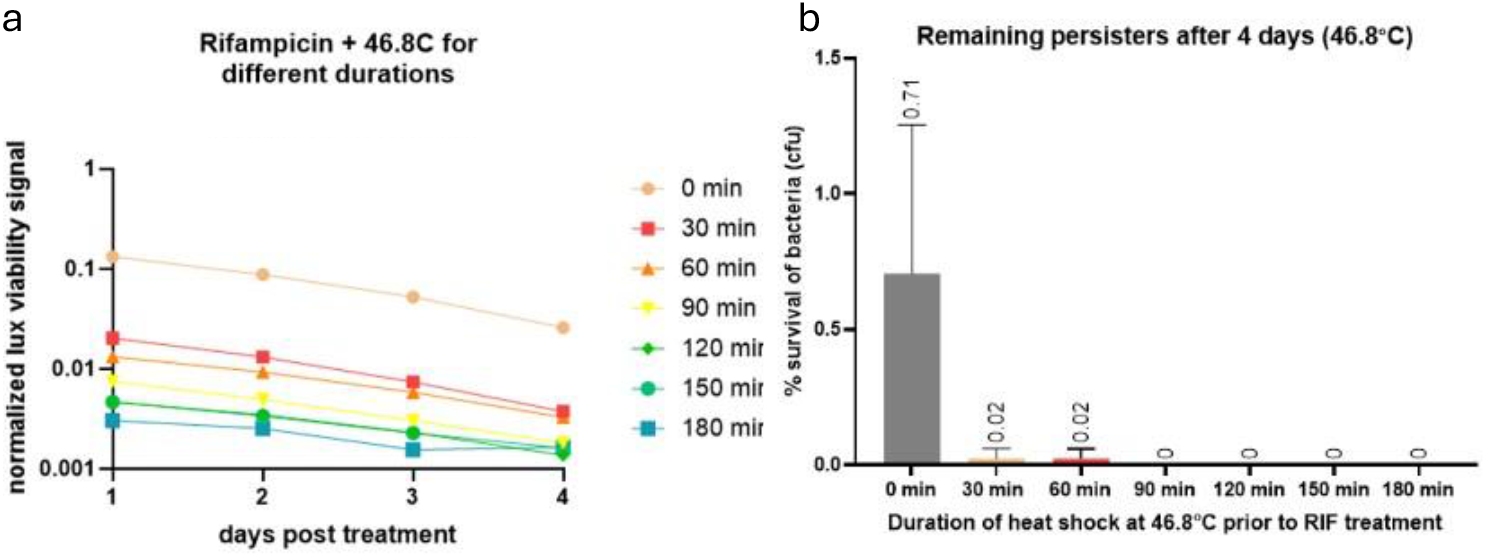
Prolonged hyperthermia enhances rifampicin-mediated killing of M. marinum. Biofilms were exposed to 46.8 °C for 30–180 min followed by 400 μg/ml rifampicin treatment. (a) In the biofilm MDK assay, the combination treatment resulted in lower antimicrobial tolerance. (b) CFU counts assessing the remaining persisters after 4 days of treatment confirmed significantly reduced bacterial survival in the combination treatment. 90 minutes hyperthermia followed by rifampicin was sufficient to clear all bacteria.

### Combining hyperthermia to streptomycin treatment increases the killing of *M. tuberculosis* H37Ra biofilms

A similar primary screen was carried out in *M. tuberculosis* H37Ra to assess whether hyperthermia could potentiate bacterial killing by antimicrobial agents. Both rifampicin and streptomycin, drugs that are bacteriocidal and effective against *M. tuberculosis* (33,34) were tested on 1-week-old and 2-week-old biofilms in combination with hyperthermia. Hyperthermic treatment alone produced results comparable to those obtained with *M. marinum*, with 60 °C sufficient to markedly reduce viability (Fig. 3a&b). For *M. tuberculosis*, rifampicin at such a high dose (400 μg/ml) used in the in vitro MDK assay, the drug alone was highly potent, and no additional effect of heat treatment could be observed (supplementary figure 1). However, when streptomycin was combined with hyperthermia, a clear potentiation was observed: in 1-week-old biofilms, temperatures of 47–51.8 °C already enhanced bacterial killing, while higher temperatures led to near-complete loss of viability in the culture (Fig 3a Heat and streptomycin). In 2-week-old biofilms, the combination resulted in a strong suppression of the viability signal at ≥51.8 °C (Fig. 3b Heat and streptomycin).

**Figure 3.**
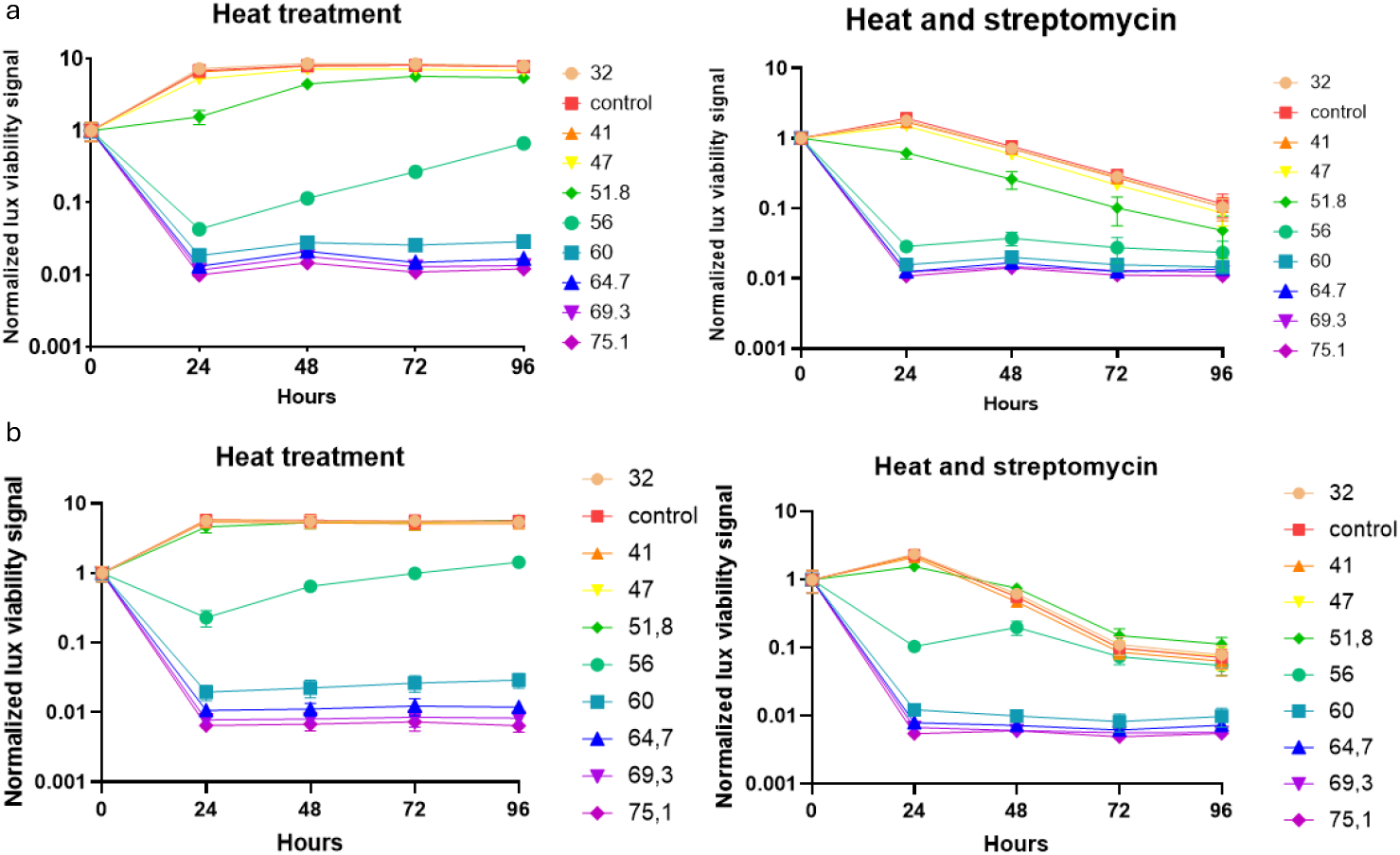
Hyperthermia potentiates streptomycin activity against M. tuberculosis biofilms. Biofilms were treated with heat (15 min across a temperature gradient) prior to streptomycin treatment. In both (a) 1-week-old biofilms and (b) 2-week-old biofilms, potentiated effects were observed at ≥51.8°C

### Combining hyperthermia with rifampicin is beneficial against *M. marinum* in *ex vivo* granulomas

To investigate whether heat treatment combined with the antimicrobial activity of rifampicin is beneficial in a more physiologically relevant setting, we analyzed *ex vivo* granulomas collected from *M. marinum*-infected zebrafish at 13 weeks post infection. Bacteria extracted from granulomas were subjected to hyperthermia (47 °C for 30 min), rifampicin (400 μg/ml) alone, or a combination of rifampicin and heat. After 24 h of the treatment application, bacterial killing was measured by cfu plating (Fig. 4). The combination treatment resulted in 4.3-fold less viable cells (P=0.0125) after 1 day of treatment suggesting that heat treatment with rifampicin combats tolerance in *in vivo* matured bacterial biofilm populations.

**Figure 4.**
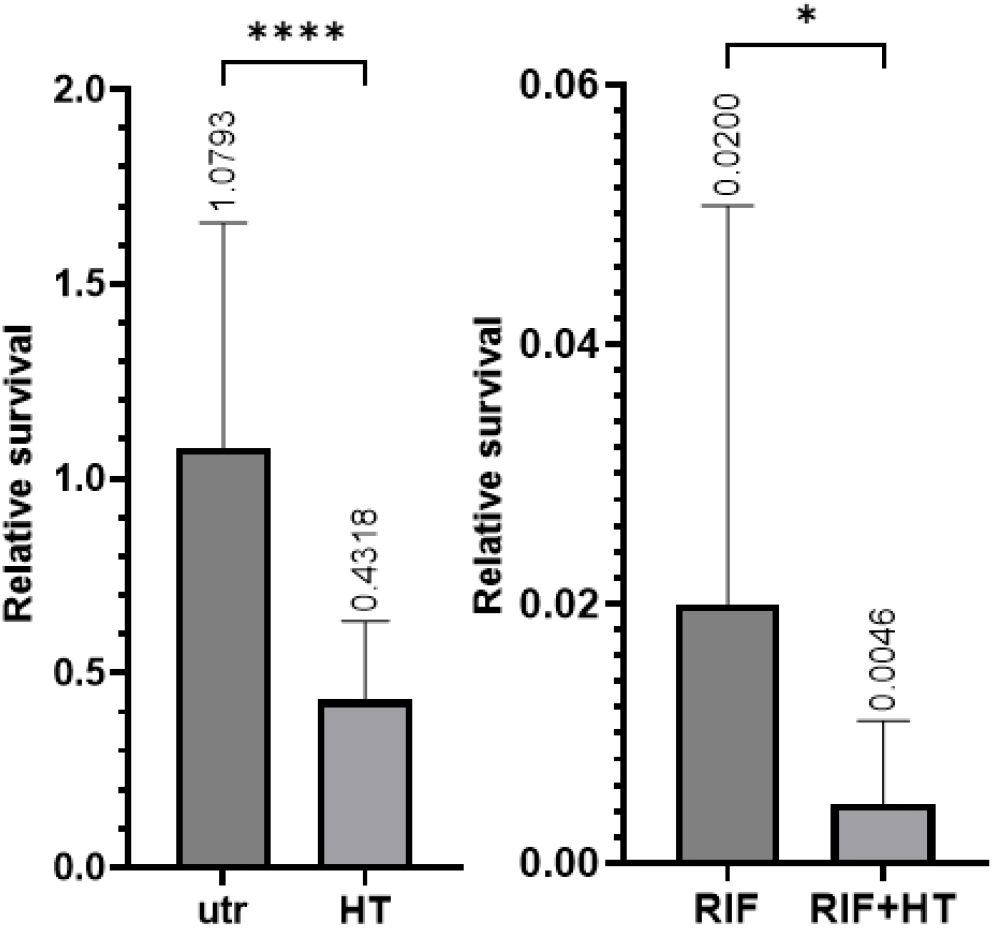
Hyperthermia combined with rifampicin potentiates bacterial killing in ex vivo granulomas. Granulomas from infected zebrafish were subjected to heat (47 °C, 30 min), rifampicin, or their combination. Combined treatment caused a stronger reduction in bacterial viability compared to either treatment alone. Friedman’s test followed by Dunn’s multiple comparisons test, error bar: mean ± SD. *p=0.0125, ****p<0.0001, n = 22.

## Discussion

Our findings demonstrate that hyperthermia significantly enhances the bacterial killing of *M. marinum* and *M. tuberculosis* biofilms, as well as in *ex vivo* granulomas of *M. marinum-* infected zebrafish. These results support the concept that hyperthermia combined with antimicrobial therapy can be beneficial against antimicrobial tolerant biofilms potentially shortening the time to cure. The beneficial effects we observed are consistent with earlier studies in non-mycobacterial pathogens. For instance, hyperthermia disrupted biofilm integrity and potentiated antimicrobial action in *P. aeruginosa* and *S. aureus* biofilms (12,13,15). Heat has been shown to induce dispersal in *P. aeruginosa* biofilms (20), and to reduce biofilm biomass while increasing planktonic cell numbers, potentially enhancing antimicrobial penetration. Importantly, our work extends this paradigm to slow-growing mycobacteria, which are notoriously tolerant due to their lipid-rich cell walls and extracellular matrix (6,31,32). The finding that heating (≈47–51 °C) is beneficial in combination with rifampicin or streptomycin suggests that thermal stress compromises these tolerance mechanisms, possibly by destabilizing membranes, altering matrix permeability, or impairing bacterial stress responses such as heat-shock protein induction.

The use of physical treatments such as heat may also have important implications for limiting the evolution of antimicrobial resistance. In contrast to drug-based therapies, which act on defined molecular targets and thus exert strong selective pressure for resistant mutants, heat causes widespread damage to proteins, membranes, and nucleic acids without favouring specific mutations (35). Mycobacteria can transiently enhance their heat tolerance through induction of heat-shock proteins, but these responses are physiological and reversible, not heritable (36). Consequently, hyperthermia is unlikely to drive stable resistance, offering a complementary approach that reduces bacterial viability without contributing to the global resistance crisis. In the context of *M. tuberculosis*, where multidrug-resistant and extensively drug-resistant strains represent a growing clinical challenge, this is particularly relevant (37). Genetic resistance in mycobacteria commonly arises through chromosomal mutations in drug targets or activation of efflux systems (38,39). These mechanisms evolve under antimicrobial pressure but would not confer protection against nonspecific physical stress. Thus, controlled hyperthermia could enhance the killing of both susceptible and resistant populations, potentially restoring antimicrobial efficacy in difficult-to-treat infections.

Importantly, this study demonstrates that the observed benefit is not limited to *in vitro* biofilms but extends to *ex vivo* granulomas of adult zebrafish, a model that more closely resembles the pathological niche of TB. Granulomas are characterized by hypoxia, nutrient limitation, and altered immune interactions that promote bacterial persistence (40). The ability of hyperthermia to potentiate bacterial killing by rifampicin even in this context highlights the translational potential of controlled hyperthermia as an adjuvant TB therapy. Since antimicrobial-tolerant biofilms in granulomas represent a major barrier to efficient eradication of persistent bacteria by antimicrobial drugs, thermal interventions that speed up the killing of tolerant bacterial populations could shorten TB treatment duration and reduce relapses.

From a clinical perspective, localized heating could be achieved using a range of emerging technologies, including magnetic nanoparticle–induced hyperthermia, alternating magnetic fields, and photothermal nanomaterials (22,23,25,26,41). The first evidence on the applicability of such methods against *M. tuberculosis in vitro* and in macrophages was recently published at the temperature of 45°C (27), in alignment with our current findings in this study. Our results suggest that hyperthermia will also be beneficial against tolerant bacteria in mycobacterial biofilms and could hence reduce the required treatment times.

Most hyperthermia-based medical applications operate within 40–44 °C to avoid thermal injury while still inducing physiological responses (42). However, short, localized exposures up to 50 °C have been reported to be tolerable in animal models and targeted cancer therapies when heat is precisely confined (43). Therefore, while the 47–51 °C range identified in this study exceeds conventional mild hyperthermia, it remains within a potentially achievable and controllable range. These findings support the feasibility of applying short-duration heating in combination with antimicrobials to overcome tolerance, provided that exposure parameters are carefully optimized for tissue safety.

## Materials and Methods

### Bacterial strains and culture conditions

*M. marinum* (ATCC927) and avirulent *M. tuberculosis* strain H37Ra (ATCC25177) were used in this study. Biofilms were cultured in 7H10 agar plate supplemented with 10% (vol/vol) of OADC (oleic acid, albumin, dextrose, catalase) enrichment (Thermo Fisher Scientific, New Hampshire, USA) and 0.5% (vol/vol) glycerol (Sigma-Aldrich, Missouri, USA). *M. marinum* was cultured at 29°C for 1 week and *M. tuberculosis* at 37°C for 3 weeks. For planktonic cultures, a clump of bacteria was transferred from the plate into Middlebrook 7H9 medium supplemented with 10% (vol/vol) ADC (Fisher Scientific, NH, USA), 0.2% (vol/vol) glycerol, and 0.2% (vol/vol) Tween 80 (Sigma-Aldrich, MO, USA). For biofilm cultures, the same medium but without glycerol and Tween80 was used, and the inoculum was adjusted to an optical density (OD600) of 0.1. The culture was then incubated at 29 °C (*M. marinum)* or at 37°C (*M. tuberculosis)* for 7/14 days.

### Primary screening of effective temperatures

#### Heat treatment screening

*M. marinum* or *M. tuberculosis* carrying the pMV306hsp+LuxG13 plasmid, a gift from Brian Robertson & Siouxsie Wiles (Addgene plasmid # 26161 ; http://n2t.net/addgene:26161 ; RRID:Addgene_26161) that expresses luciferase, were cultured in biofilm-forming conditions at the volume of 10mls. Cultures were homogenized by pipetting and divided into sterile PCR plates. Heating was carried out using Bio-Rad C1000 Thermal Cycler. Bacteria were heated for 15 minutes at the temperature of 32.0, 36.5, 41.1, 47.0, 51.8, 56.0, 60.0, 64.7, 69.3 or 75.1°C. After the heating, samples were transferred to sterile white 96-well plates at the volume of 192μl per well, 6 samples per temperature. To half of the *M. marinum* samples, rifampicin (400 μg/ml) was added. In the case of *M. tuberculosis* either rifampicin (400 μg/ml) or streptomycin (800 μg/ml) was used.

#### Biofilm minimum duration for killing (MDK) assay

Bioluminescent strains of *M. marinum* and *M. tuberculosis* transformed with a plasmid containing a constitutive bioluminescence cassette (pMV306hsp+LuxG13) (Addgene plasmid # 26161; http://n2t.net/addgene:26161; RRID:Addgene_ were used. The plasmid allows the constitutive production of luciferases and its substrates and repetitive measurements of the viability signal. Bacteria were cultured to biofilms as described above. Bioluminescence was measured over time using an EnVision® multilabel plate reader (PerkinElmer). For each well, five readings of 1 second each were recorded, and the mean value was used for subsequent analysis. Assays were performed with 3–6 biological replicates. For each time point, bioluminescence values were normalized to the initial measurement. Using Office Excel and GraphPad Prism 10, the values were plotted to create lux-based time-kill curves and to calculate AUC-values.

### Screening 30-180-minute treatments

For assessing lower temperatures with longer treatment times, *M. marinum* was cultured, homogenized and divided into a PCR plate as in the primary screen. Heating temperatures were set to 45.6, 46.8, 49.4 or 51.3°C and the samples were heated for 30, 60, 90 or 120 minutes. The samples were divided into white 96-well plate and measured as before with biofilm MDK assay. After 4 days at the endpoint, CFUs were determined from the samples. The samples were homogenized, washed 3 times with 200μls of sterile PBS and plated to determine the number of viable bacteria. Colonies were counted after 6 days of culture.

### Zebrafish housing and ethics statement

Female zebrafish (4–12 months old) obtained from Tampere Zebrafish Core Facility were used for the in vivo experiment. Fish were maintained in a flow-through housing system with a 14 h light/10 h dark cycle and fed GEMMA Micro 500 (Skretting, USA) daily. Water conditions were kept at pH 7.6 and conductivity 1000 μS using sea salt and NaHCO3, and the temperature was maintained at 28 °C. Fish were housed in groups of 10–25 individuals. All zebrafish husbandry and experimental procedures were approved by the Animal Experiment Board in Finland (licenses ESAVI/14286/2022 and ESAVI/10079/04) and conducted in compliance with EU Directive 2010/63/EU on the protection of animals used for scientific purposes and the Finnish Act on Animal Experimentation (62/2006).

### Zebrafish infections

Planktonic *M. marinum* was grown to an OD600 of 0.5, with one intermediate dilution performed 2 days before fish infection. A 1 ml of liquid culture was centrifuged (3 min, 10,000 × g), the supernatant discarded, and the pellet resuspended and diluted in 1× PBS containing 0.3 mg/mL phenol red. Dilutions were made according to the predetermined OD600–CFU/μl curve to achieve a final concentration of 7 CFU/μl. The bacterial suspension was passed three times through a 27 G needle. Fish were anesthetized in 0.02% 3 aminobenzoic acid ethyl ester and injected intracoelomically with 5 μl using an Omnican 100, 30 G insulin needle (Braun, Melsungen, Germany). Multiple samples were plated on 7H10 agar during the infection procedure to confirm the infection dose. After *M. marinum* infection, humane endpoint criteria approved by the national ethical board were followed. Fish exhibiting any of the following signs: lack of response to touch, abnormal swimming, gasping, visible swelling, wasting, or loss of scales, were euthanized with an overdose of 3 aminobenzoic acid ethyl ester.

### Ex vivo hyperthermia

Granulomas were collected from multiple infected fish, with 20 granulomas pooled per sample tube from a single individual fish to serve as one biological replicate. Each tube was supplemented with 400 μL of sterile PBS. Samples were homogenized by adding beads and processing them using a bead beater (FastPrep-24 5G, 6.5 m/s for 2×40-second cycles with a 60-second pause). Serial dilutions of the homogenates were plated to determine baseline bacterial loads. Following homogenization, samples were transferred to 2 mL Eppendorf tubes and brought to a final volume of 1.8 mL by adding an additional 1,400 μL of sterile PBS. Hygromycin (50 μg/mL, InvivoGen) was added to each tube to eliminate potential contamination. Aliquots of 192 μL were taken from each sample for the non-heated control condition. The remaining samples were subjected to heat shock by incubation in a heat block at 47 °C for 30 minutes. Post-treatment, 192 μL aliquots were distributed into wells of a 96-well plate. Rifampicin (8 μL per well) was added to achieve a final concentration of 400 μg/mL; sterile water was added to control wells. At designated time points (0- and 1-days post-treatment), serial dilutions were plated to assess bacterial viability.

### Statistical analysis

Microsoft Excel and GrapPad Prism 10 were used for data processing. All visualizations were generated in Prism 10. One-way ANOVA with Dunnett’s multiple comparisons test was used to assess statistical significance.

## Supplementary figures

**Supplementary figure 1.**
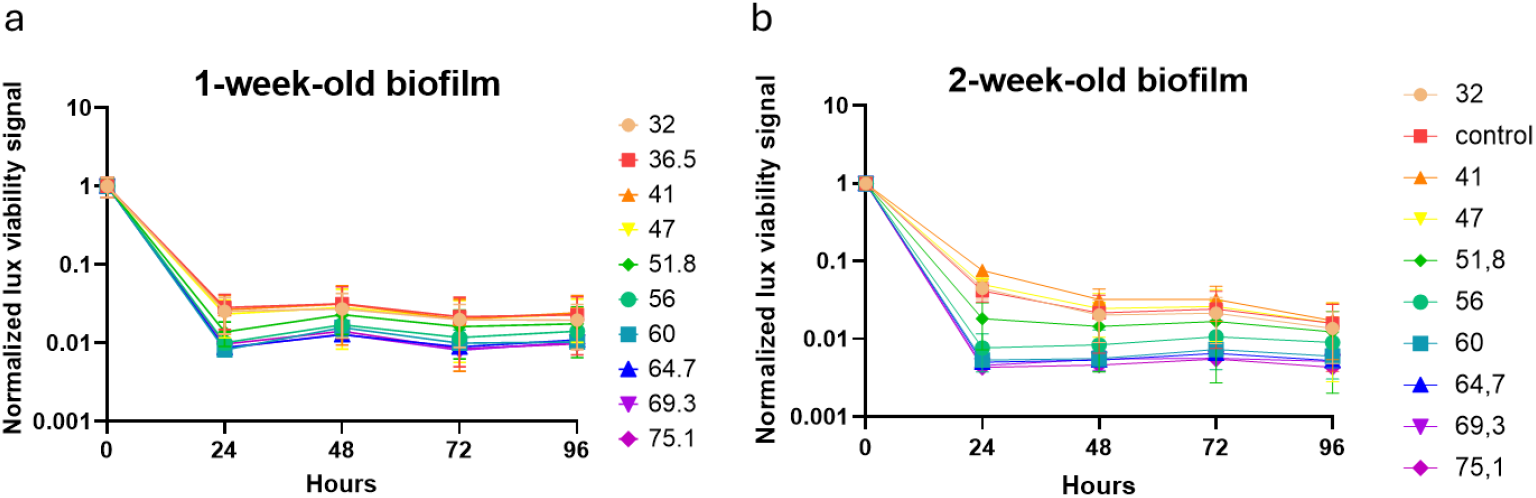
Rifampicin alone was highly potent in both 1- and 2-week-old biofilms. Biofilms were exposed to heat for 15 minutes across a temperature gradient prior to rifampicin treatment. No additional enhancement by heat was detected in either (a) 1-week-old or (b) 2-week-old biofilms, likely due to the high concentration of rifampicin (400μg/ml).

## Acknowledgements

The authors acknowledge the Biocenter Finland-funded TUNIFish Tampere Zebrafish Laboratory for the service. Mataleena Parikka research group is part of the Research Center for Vaccine Development and Immunology (VACCIM) at Tampere University. We acknowledge Hannaleena Piippo and Leena Mäkinen for their technical assistance in this work.

## Notes

### Competing Interest Statement

A patent application related to the methodology used in this article has been submitted by Tampere University in July 2025. MH and MP are original inventors in this patent application.

